# Cryo-EM Structure of the 2019-nCoV Spike in the Prefusion Conformation

**DOI:** 10.1101/2020.02.11.944462

**Authors:** Daniel Wrapp, Nianshuang Wang, Kizzmekia S. Corbett, Jory A. Goldsmith, Ching-Lin Hsieh, Olubukola Abiona, Barney S. Graham, Jason S. McLellan

**Author notes:** These authors contributed equally (Daniel Wrapp, Nianshuang Wang). Correspondence to (J.S.M.).

## Abstract

The outbreak of a novel betacoronavirus (2019-nCov) represents a pandemic threat that has been declared a public health emergency of international concern. The CoV spike (S) glycoprotein is a key target for urgently needed vaccines, therapeutic antibodies, and diagnostics. To facilitate medical countermeasure (MCM) development we determined a 3.5 Å-resolution cryo-EM structure of the 2019-nCoV S trimer in the prefusion conformation. The predominant state of the trimer has one of the three receptor-binding domains (RBDs) rotated up in a receptor-accessible conformation. We also show biophysical and structural evidence that the 2019-nCoV S binds ACE2 with higher affinity than SARS-CoV S. Additionally we tested several published SARS-CoV RBD-specific monoclonal antibodies and found that they do not have appreciable binding to nCoV-2019 S, suggesting antibody cross-reactivity may be limited between the two virus RBDs. The atomic-resolution structure of 2019-nCoV S should enable rapid development and evaluation of MCMs to address the ongoing public health crisis.

---

The novel coronavirus 2019-nCoV has recently emerged as a human pathogen in the city of Wuhan in China’s Hubei province, causing fever, severe respiratory illness and pneumonia (*1, 2*). According to the World Health Organization on February 10^th^, 2020, there have been over 40,000 confirmed cases globally, leading to at least 900 deaths. The new pathogen was rapidly shown to be a novel member of the betacoronavirus genus that is closely related to several bat coronaviruses as well as severe acute respiratory syndrome coronavirus (SARS-CoV) (*3, 4*). Compared to SARS-CoV, 2019-nCoV appears to be more readily transmitted from human-to-human, spreading to multiple continents and leading to the WHO declaration of a Public Health Emergency of International Concern (PHEIC) (*1, 5, 6*).

2019-nCoV makes use of a densely glycosylated, homotrimeric class I fusion spike (S) protein to gain entry into host cells. The S protein exists in a metastable prefusion conformation that undergoes a dramatic structural rearrangement to fuse the viral membrane with the host cell membrane (*7, 8*). This process is triggered by binding of the S1 subunit to a host-cell receptor, which destabilizes the prefusion trimer, resulting in shedding of the S1 subunit and transition of the S2 subunit to a highly stable postfusion conformation (*9*). In order to engage a host-cell receptor, the receptor-binding domain (RBD) of S1 undergoes hinge-like conformational movements that transiently hide or expose the determinants of receptor binding. These two states are referred to as the “down” conformation and the “up” conformation, where “down” corresponds to the receptor-inaccessible state and “up” corresponds to the receptor-accessible state, which is thought to be less stable (*10-13*). Due to the indispensable function of the S protein it represents a vulnerable target for antibody-mediated neutralization, and characterization of the prefusion S structure would provide atomic-level information to guide vaccine design and development.

Based on the reported genome sequence of 2019-nCoV(*4*), we expressed ectodomain residues 1−1208 of 2019-nCoV S (**Figure 1A, Supplementary Figure 1**), adding two stabilizing proline mutations in the C-terminal S2 fusion machinery based on a previous stabilization strategy which proved highly effective for betacoronavirus S proteins (*11, 14*). We obtained roughly 0.5 mg/L of the recombinant prefusion-stabilized S ectodomain from FreeStyle 293 cells, and the protein was purified to homogeneity by affinity chromatography and size-exclusion chromatography (**Supplementary Figure 1**). Cryo-EM grids were prepared using this purified, fully glycosylated S protein and preliminary screening revealed a high particle density with little aggregation near the edges of the holes.

**Figure 1.**
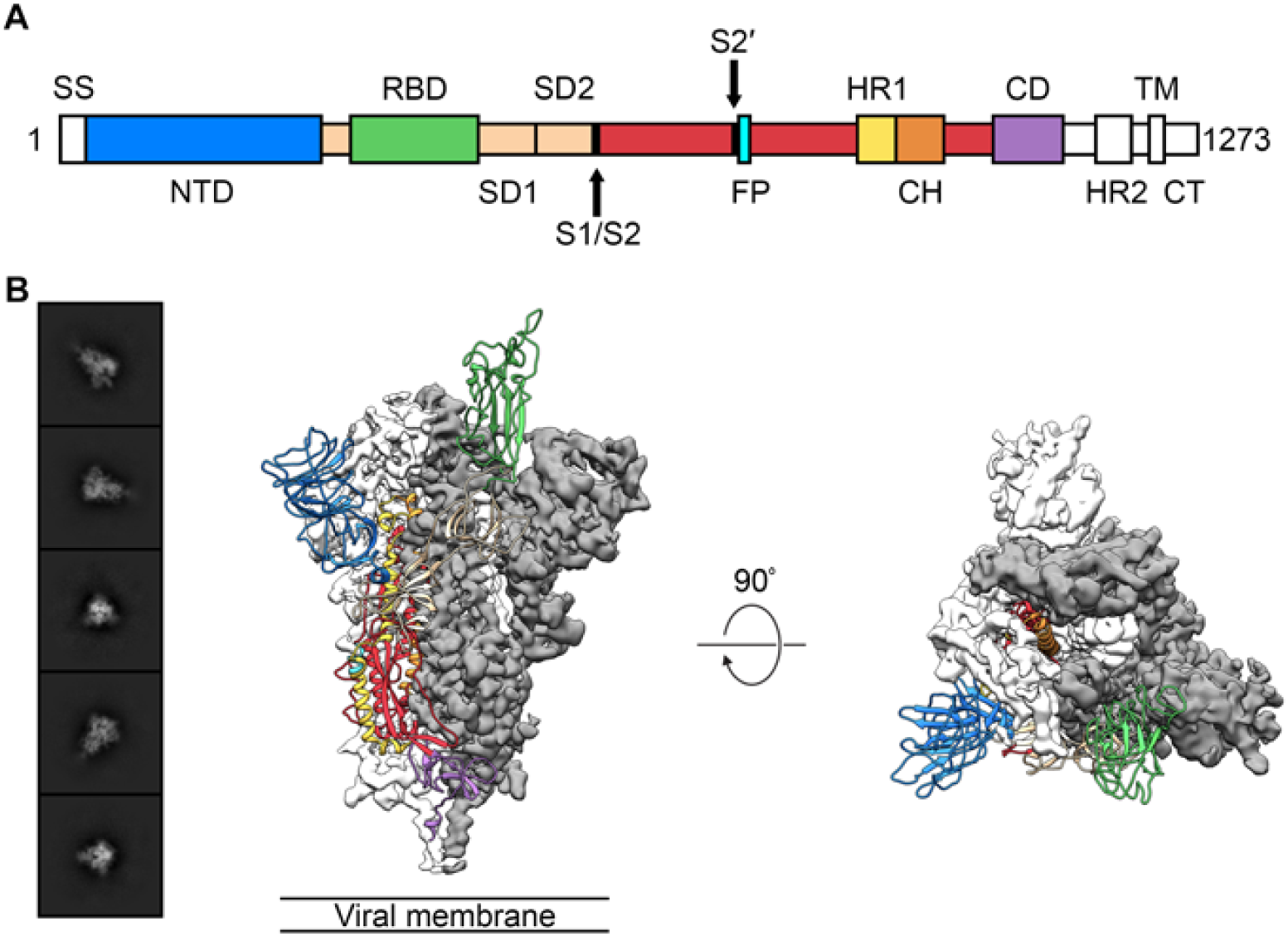
Structure of 2019-nCoV S in the prefusion conformation. (**A**) Schematic of 2019-nCoV S primary structure, colored by domain. Domains that were excluded from the ectodomain expression construct or could not be visualized in the final map are colored white. SS= signal sequence, NTD= N-terminal domain, RBD= receptor-binding domain, SD1= subdomain 1, SD2= subdomain 2, S1/S2= S1/S2 protease cleavage site, S2′= S2′ protease cleavage site, FP= fusion peptide, HR1= heptad repeat 1, CH= central helix, CD= connector domain, HR2= heptad repeat 2, TM= transmembrane domain, CT= cytoplasmic tail. Arrows denote protease cleavage sites. (**B**) Select 2D class averages of the particles that were used to calculate the 2019-nCoV S reconstruction (*left*). Side and top views of the prefusion structure of the 2019-nCoV S protein with a single RBD in the “up” conformation (*right*). The two RBD “down” protomers are shown as cryo-EM density in either white or gray and the RBD “up” protomer is shown in ribbons, colored corresponding to the schematic in **Fig 1A**.

After collecting and processing 3,207 micrograph movies, we obtained a 3.5 Å-resolution 3D reconstruction of an asymmetrical trimer in which a single RBD was observed in the “up” conformation. (**Figure 1B, Supplementary Figure 2**). Due to the small size of the RBD (∼21 kDa), the asymmetry of this conformation was not readily apparent until *ab initio* 3D reconstruction and 3D classification were performed (**Figure 1B, Supplementary Figure 3**). By using the 3D variability feature in cryoSPARC v2 (*15*), we were able to observe breathing of the S1 subunits as the RBD underwent a hinge-like movement, which likely contributed to the relatively poor local resolution of S1 compared to the more stable S2 subunit (**Supplementary Movies 1 and 2**). This seemingly stochastic RBD movement has been captured during structural characterization of the closely related betacoronaviruses SARS-CoV and MERS-CoV, as well as the more distantly related alphacoronavirus porcine epidemic diarrhea virus (PEDV)(*10, 11, 13, 16*). The observation of this phenomenon in 2019-nCoV S suggests that it shares the same mechanism of triggering that is thought to be conserved among the *Coronaviridae*, wherein receptor-binding to exposed RBDs leads to an unstable 3 RBD-up conformation that results in shedding of S1 and refolding of S2 (*11, 12*).

Because the S2 subunit appeared to be a symmetric trimer, we performed a 3D refinement imposing C3 symmetry, resulting in a 3.2 Å-resolution map, with excellent density for the S2 subunit. Using both maps we built the vast majority of the 2019-nCoV S ectodomain, including glycans at 44 of the 66 *N*-linked glycosylation sites per trimer (**Supplementary Figure 4**). Our final model spans S residues 27–1146, with several flexible loops omitted. Like all previously reported coronavirus S ectodomain structures, the density for 2019-nCoV S begins to fade after the connector domain (CD), reflecting the flexibility of the heptad repeat 2 (HR2) domain in the prefusion conformation (**Supplementary Figure 4A**) (*13, 16-18*).

The overall structure of 2019-nCoV S resembles that of SARS-CoV S, with a root mean square deviation (RMSD) of 3.8 Å over 959 Cα atoms. The largest discrepancy between these two structures is a conformational difference between the positions of the RBDs in their respective “down” conformations (**Figure 2A**). Whereas the SARS-CoV RBD in the “down” conformation packs tightly against the N-terminal domain (NTD) of the neighboring protomer, the 2019-nCoV RBD in the “down” conformation is angled closer to the central cavity of the homotrimer. Despite this observed conformational difference, when the individual structural domains of 2019-nCoV S are aligned to their counterparts from SARS-CoV S, they reflect the high degree of structural homology between the two proteins, with the NTDs, RBDs, subdomains 1 and 2 (SD1 and SD2) and S2 subunits yielding RMSD values of 2.6 Å, 3.0 Å, 2.7 Å and 2.0 Å, respectively (**Figure 2B**).

**Figure 2.**
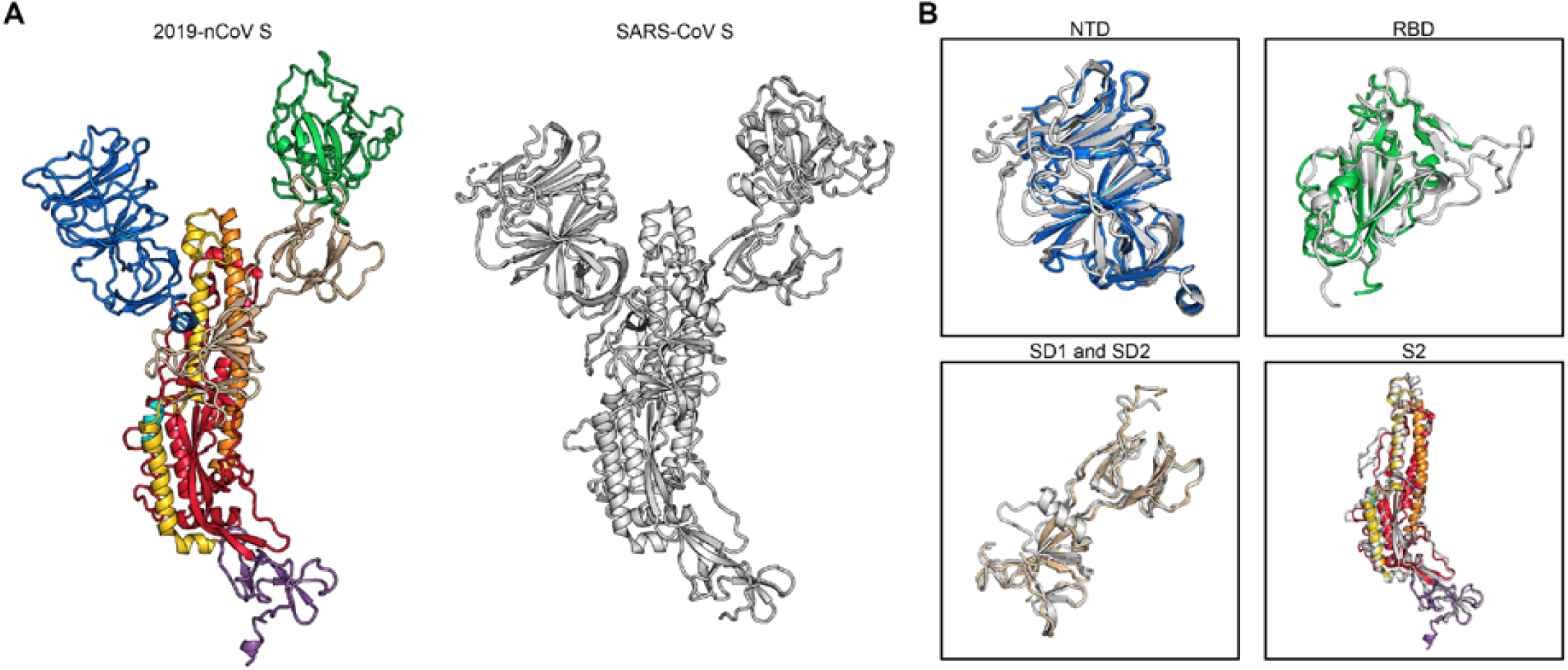
Structural comparison between 2019-nCoV S and SARS-CoV S. (**A**) A single RBD “down” monomer of 2019-nCoV S is shown in ribbons, colored according to **Figure 1**. A monomer of SARS-CoV S is also shown in ribbons, colored white (PDB ID: 6CRZ). (**B**) The following structural domains from 2019-nCoV S have been aligned to their counterparts from SARS-CoV S; NTD (*top left*), RBD (*top right*), SD1 and SD2, (*bottom left*) and S2 (*bottom right*).

2019-nCoV S shares roughly 96% sequence identity with the S protein from the bat coronavirus RaTG13, with the most notable variation arising from an insertion in the S1/S2 protease cleavage site that results in an “RRAR” furin recognition site in 2019-nCoV, rather than the single arginine in SARS-CoV (**Supplementary Figure 5**) (*19-22*). A similar phenomenon has been observed for influenza viruses, where amino acid insertions that create a polybasic furin site in a related position in influenza hemagglutinin proteins are often found in highly virulent avian and human influenza viruses (*23*). In addition to this insertion of residues in the S1/S2 junction, 29 variant residues exist between 2019-nCoV S and RaTG13 S, with 17 of these positions mapping to the RBD (**Supplementary Figures 5 and 6**). We also analyzed the 61 available 2019-nCoV S sequences in GISAID and found that there were only 9 amino acid substitutions among all deposited sequences. Most of these substitutions are relatively conservative and they are not expected to have a dramatic effect on the structure or function of the 2019-nCoV S protein (**Supplementary Figure 6**).

Recent reports demonstrating that 2019-nCoV S and SARS-CoV S share the same functional host-cell receptor—angiotensin-converting enzyme 2 (ACE2) (*21, 24-26*)—prompted us to quantify the kinetics mediating this interaction via surface plasmon resonance (SPR). Surprisingly, ACE2 bound to 2019-nCoV S ectodomain with ∼15 nM affinity, which is approximately 10- to 20-fold higher affinity than ACE2 binding to SARS-CoV S (**Figure 3A, Supplementary Figure 7**) (*14*). We also formed a complex of ACE2 bound to the 2019-nCoV S ectodomain and observed it by negative-stain EM, where it strongly resembled the complex formed between SARS-CoV S and ACE2, which has been observed at high-resolution by cryo-EM (**Figure 3B**) (*14, 27*). The high affinity of 2019-nCoV S for human ACE2 may contribute to the apparent ease with which 2019-nCoV can spread from human-to-human(*1*), however additional studies are needed to investigate this possibility.

**Figure 3.**
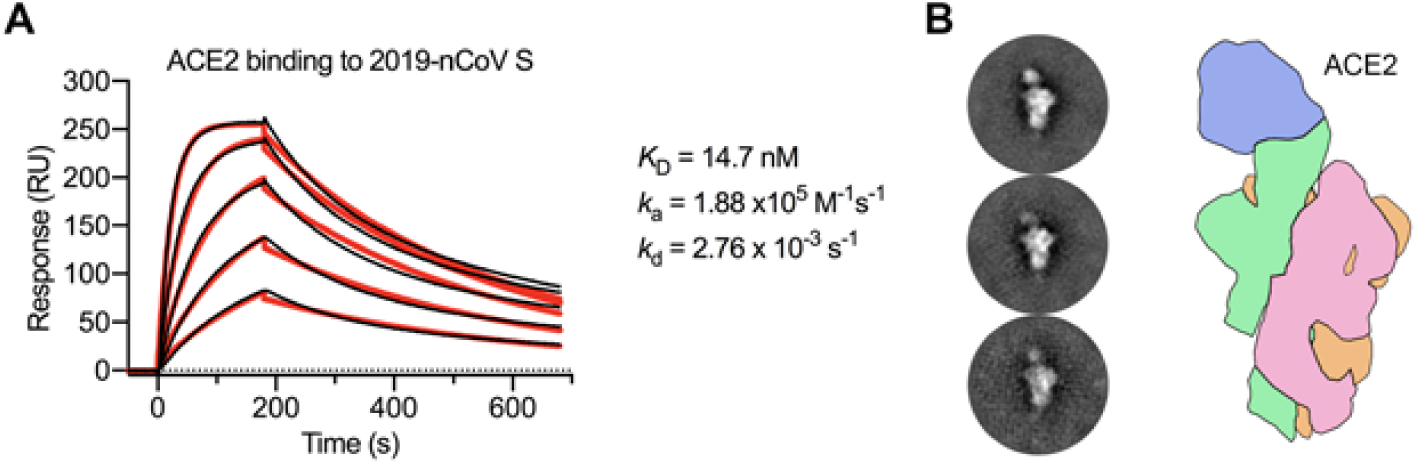
2019-nCoV S binds human ACE2 with high affinity. (**A**) SPR sensorgram showing the binding kinetics for human ACE2 and immobilized 2019-nCoV S. Data are shown as black lines and the best fit of the data to a 1:1 binding model is shown in red. (**B**) Negative-stain EM 2D class averages of 2019-nCoV S bound by ACE2. Averages have been rotated so that ACE2 is positioned above the 2019-nCoV S protein with respect to the viral membrane. A cartoon depicting the ACE2-bound 2019-nCoV S protein is shown (*right*) with ACE2 in blue and S protein monomers colored tan, pink and green.

The overall structural homology and shared receptor usage between SARS-CoV S and 2019-nCoV S prompted us to test published SARS-CoV RBD-directed monoclonal antibodies (mAbs) for cross-reactivity to the 2019-nCoV RBD (**Figure 4A**). A 2019-nCoV RBD-SD1 fragment (S residues 319–591) was recombinantly expressed, and appropriate folding of this construct was validated by measuring ACE2 binding using biolayer interferometry (BLI) (**Figure 4B**). Cross-reactivity of the SARS-CoV RBD-directed mAbs S230, m396 and 80R was then evaluated by BLI (*12, 28-30*). Despite the relatively high degree of structural homology between the 2019-nCoV RBD and the SARS-CoV RBD, no binding to the 2019-nCoV RBD could be detected for any of the three mAbs at the concentration tested (1 μM) (**Figure 4C, Supplementary Figure 8**). Although the epitopes of these three antibodies represent a relatively small percentage of the surface area of the 2019-nCoV RBD, the lack of observed binding suggests that SARS-directed mAbs will not necessarily be cross-reactive and that future antibody isolation and therapeutic design efforts will benefit from using 2019-nCoV S proteins as probes.

**Figure 4.**
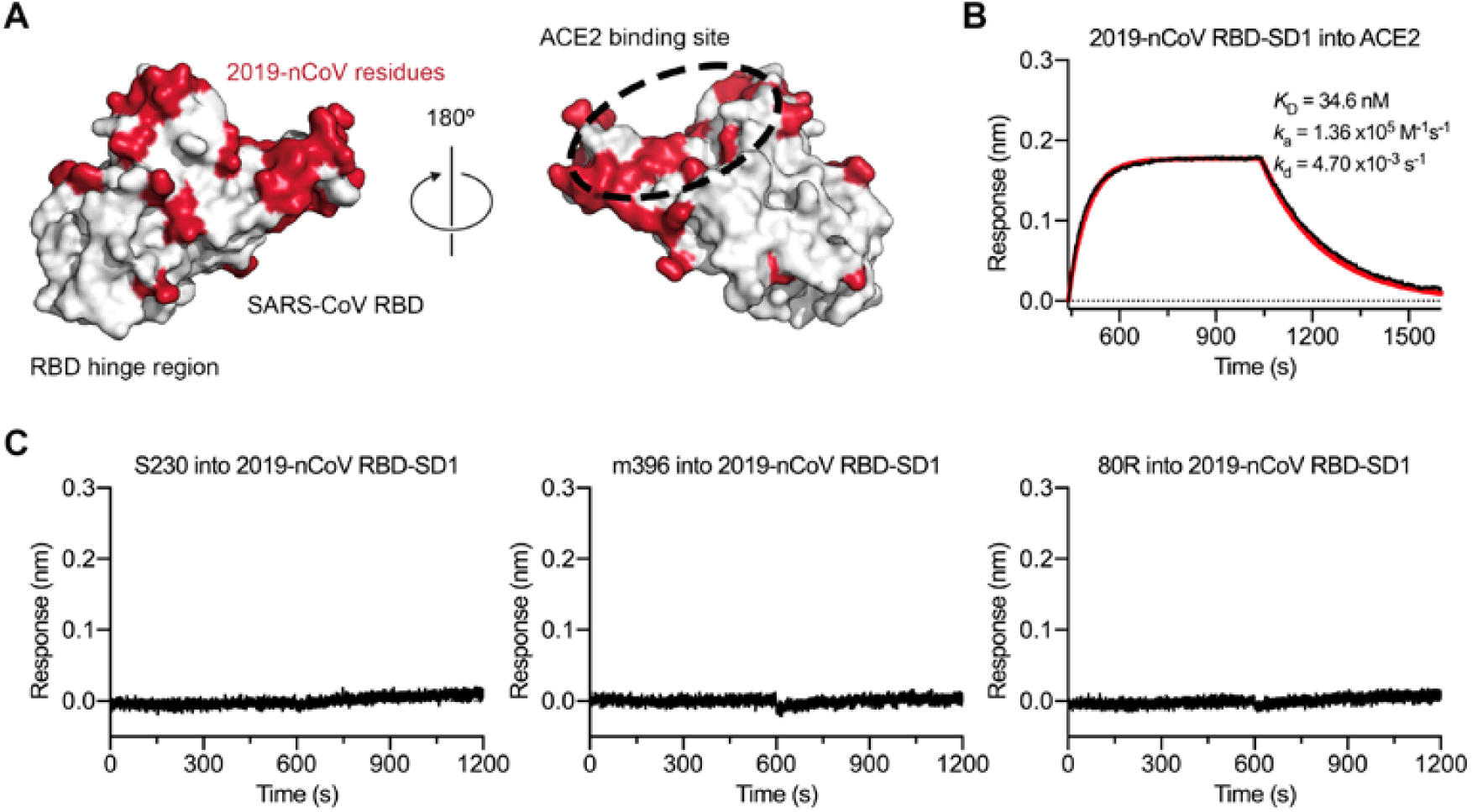
Antigenicity of the 2019-nCoV RBD. (**A**) The SARS-CoV RBD is shown as a white molecular surface (PDB ID: 2AJF), with residues that vary in the 2019-nCoV RBD colored red. The ACE2 binding site is outlined with a black dotted line. (**B**) A biolayer interferometry sensorgram that shows binding to ACE2 by the 2019-nCoV RBD-SD1. Binding data are shown as a black line and the best fit of the data to a 1:1 binding model is shown in red. (**C**) Biolayer interferometry to measure cross-reactivity of the SARS-CoV RBD-directed antibodies S230, m396 and 80R. Sensortips with immobilized antibodies were dipped into wells containing 2019-nCoV RBD-SD1 and the resulting data are shown as a black line.

The rapid global spread of 2019-nCoV, prompting the PHEIC declaration by WHO signals the urgent need for coronavirus vaccines and therapeutics. Knowing the atomic-level structure of the spike will support precision vaccine design and discovery of antivirals, facilitating medical countermeasure development.

## Materials and Methods

### Protein expression and purification

To express the prefusion S ectodomain, a gene encoding residues 1−1208 of 2019-nCoV S (GenBank: MN908947) with proline substitutions at residues 986 and 987, a “GSAS” substitution at the furin cleavage site (residues 682–685), a C-terminal T4 fibritin trimerization motif, an HRV3C protease cleavage site, a TwinStrepTag and an 8XHisTag was synthesized and cloned into the mammalian expression vector pαH. To express the 2019-nCoV RBD-SD1, residues 319−591 of 2019-nCoV S were cloned upstream of a C-terminal HRV3C protease cleavage site, a monomeric Fc tag and an 8XHisTag. Similarly, to express the SARS-CoV RBD-SD1, residues 306−577 of SARS-CoV S (Urbani strain) were cloned upstream of a C-terminal HRV3C protease cleavage site, a monomeric Fc tag and an 8XHisTag. Lastly, a plasmid encoding residues 1−615 of human ACE2 with a C-terminal HRV3C protease cleavage site, a TwinStrepTag and an 8XHisTag was generated.

These expression vectors were used to transiently transfect FreeStyle293F cells (Thermo Fischer) using polyethylenimine. Protein was purified from filtered cell supernatants using either StrepTactin resin (IBA) or Protein A resin (Pierce) before being subjected to additional purification by size-exclusion chromatography using either a Superose 6 10/300 column (GE Healthcare) or a Superdex 200 10/300 Increase column (GE Healthcare) in 2 mM Tris pH 8.0, 200 mM NaCl and 0.02% NaN_3_. ACE2 and the 2019-nCoV RBD-SD1 were incubated with 10% (wt/wt) HRV3C protease for 2 hours at room temperature. Cleaved protein was then passed over either NiNTA resin (ACE2) or Protein A and NiNTA resins (2019-nCoV RBD) to remove cleaved tags and His-tagged protease before being run over a Superdex 200 10/300 Increase column in 2 mM Tris pH 8.0, 200 mM NaCl and 0.02% NaN_3_.

Plasmids encoding the heavy and light chains of S230, 80R and m396 IgG were transiently transfected into Expi293 (Thermo Fischer) using polyethylenimine. Antibodies were purified from cell supernatants using Protein A resin before being used for biolayer interferometry.

### Cryo-EM sample preparation and data collection

Purified 2019-nCoV S was diluted to a concentration of 0.35 mg/mL in 2 mM Tris pH 8.0, 200 mM NaCl and 0.02% NaN_3_. 3 uL of protein was deposited on a CF-1.2/1.3 grid that had been plasma cleaned for 30 seconds in a Solarus 950 plasma cleaner (Gatan) with a 4:1 ratio of O_2_/H_2_. Excess protein was blotted away for 6 seconds before being plunge frozen into liquid ethane using a Vitrobot Mark IV (Thermo Scientific). Frozen grids were imaged in a Titan Krios (Thermo Scientific) equipped with a K3 detector (Gatan). Movies were collected using Leginon *(31)* at a magnification of x22,500, corresponding to a calibrated pixel size of 1.047 Å/pixel. A full description of the cryo-EM data collection parameters can be found in **Supplementary Table 1**.

### Cryo-EM data processing

Motion correction, CTF-estimation and non-templated particle picking were performed in Warp (*32*). Extracted particles were imported into cryoSPARC v2.12.4 (*15*) for 2D classification, 3D classification and non-uniform 3D refinement. The C1 RBD “up” reconstruction was sharpened in cryoSPARC, and the 3D reconstruction with C3 symmetry was subjected to local B-factor sharpening using LocalDeBlur (*33*). Models were built in Coot, before being iteratively refined in both Phenix and ISOLDE (*34-36*). Some of the data processing and refinement software was curated by SBGrid (*37*). The full cryo-EM data processing workflow is described in **Supplementary Figure 3** and the model refinement statistics can be found in **Supplementary Table 1**.

### Surface plasmon resonance

His-tagged 2019-nCoV S was immobilized to an NiNTA sensorchip (GE Healthcare) to a level of ∼800 response units (RUs) using a Biacore X100 (GE Healthcare) and a running buffer composed of 10 mM HEPES pH 8.0, 150 mM NaCl and 0.05% Tween 20. Serial dilutions of purified and untagged ACE2 were injected ranging in concentration from 250 to 15.6 nM. The resulting data were fit to a 1:1 binding model using Biacore Evaluation Software (GE Healthcare). His-tagged SARS-CoV RBD-SD1 was immobilized to an NiNTA sensorchip to a level of ∼350 RUs using a Biacore X100 and the same running buffer listed above. Serial dilutions of purified and untagged ACE2 were injected ranging in concentration from 500 to 31.3 nM. The resulting data were fit to a 1:1 binding model using Biacore Evaluation Software.

### Negative stain EM

Purified 2019-nCoV S was diluted to a concentration of 0.032 mg/mL in 2 mM Tris pH 8.0, 200 mM NaCl and 0.02% NaN_3_. Diluted S protein was mixed with a 1.5-fold molar excess of ACE2 and the mixture was incubated on ice for 1 minute before 4.8 uL of the protein mixture was deposited on a CF400-Cu grid (Electron Microscopy Sciences) before being stained with methylamine tungstate (Nanoprobes). This grid was imaged in an FEI Talos TEM (Thermo Scientific) equipped with a Ceta 16M detector. Micrographs were collected manually using TIA v4.14 software at a magnification of x92,000, corresponding to a pixel size of 1.63 Å/pixel. CTF estimation, particle picking and 2D class averaging were performed in *cis*TEM (*38*).

### Biolayer interferometry

Fc-tagged 2019-nCoV RBD-SD1 was immobilized to an anti-human capture (AHC) sensortip (FortéBio) using an Octet RED96e (FortéBio). The sensortip was then dipped into 100 nM ACE2 to measure association before being dipped into a well containing only running buffer composed of 10 mM HEPES pH 7.5, 150 mM NaCl, 3 mM EDTA, 0.05% Tween 20 and 1 mg/mL bovine serum albumin to measure dissociation. Data were reference subtracted and fit to a 1:1 binding model using Octet Data Analysis Software v11.1 (FortéBio).

S230, 80R and m396 IgGs were immobilized to AHC sensortips to a response level of ∼0.8 nm and dipped into wells containing 1 μM untagged 2019-nCoV RBD-SD1 before being dipped into wells containing only running buffer to measure dissociation. Data were reference-subtracted and aligned to a baseline after IgG capture using Octet Data Analysis software v11.1. An analogous experiment was performed under identical conditions by dipping AHC sensor tips loaded with S230, 80R or m396 IgG into untagged SARS-CoV RBD-SD1. Data were reference-subtracted, aligned to a baseline after IgG capture and fit to a 1:1 binding model using Octet Data Analysis software v11.1.

## Acknowledgments

We thank Dr. John Ludes-Meyers for assistance with cell transfection and the rest of the members of the McLellan laboratory for critical reading of the manuscript. We would also like to thank Dr. Aguang Dai from the Sauer Structural Biology Laboratory at the University of Texas at Austin for his assistance with microscope alignment. This work was supported by an NIH/NIAID grant awarded to J.S.M. (R01-AI127521). The Sauer Structural Biology Laboratory is supported by the University of Texas College of Natural Sciences and by award RR160023 of the Cancer Prevention and Research Institute of Texas (CPRIT).

---

**Supplementary Figure 1.**
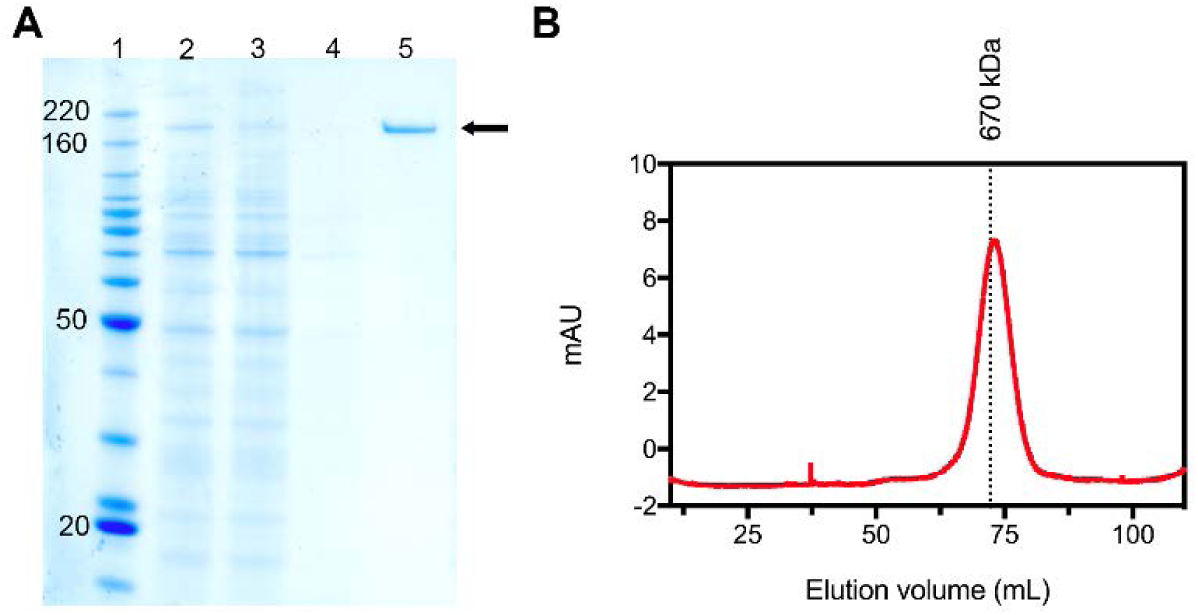
2019-nCoV S expression and purification. (**A**) SDS-PAGE analysis of the 2019-nCoV S protein. Lane 1: molecular weight ladder, with relevant bands labeled in kilodaltons (*left*); lane 2: filtered supernatant from transfected cells; lane 3: supernatant after passing through StrepTactin resin; lane 4: wash of StrepTactin resin; lane 5: elution from StrepTactin resin. The band corresponding to 2019-nCoV S is denoted with a black arrow. (**B**) Size-exclusion chromatogram of the affinity-purified 2019-nCoV S protein. Data from a Superose 6 10/300 column are shown in red. The elution volume of a 670 kilodalton molecular weight standard is shown as a black dotted line.

**Supplementary Figure 2.**
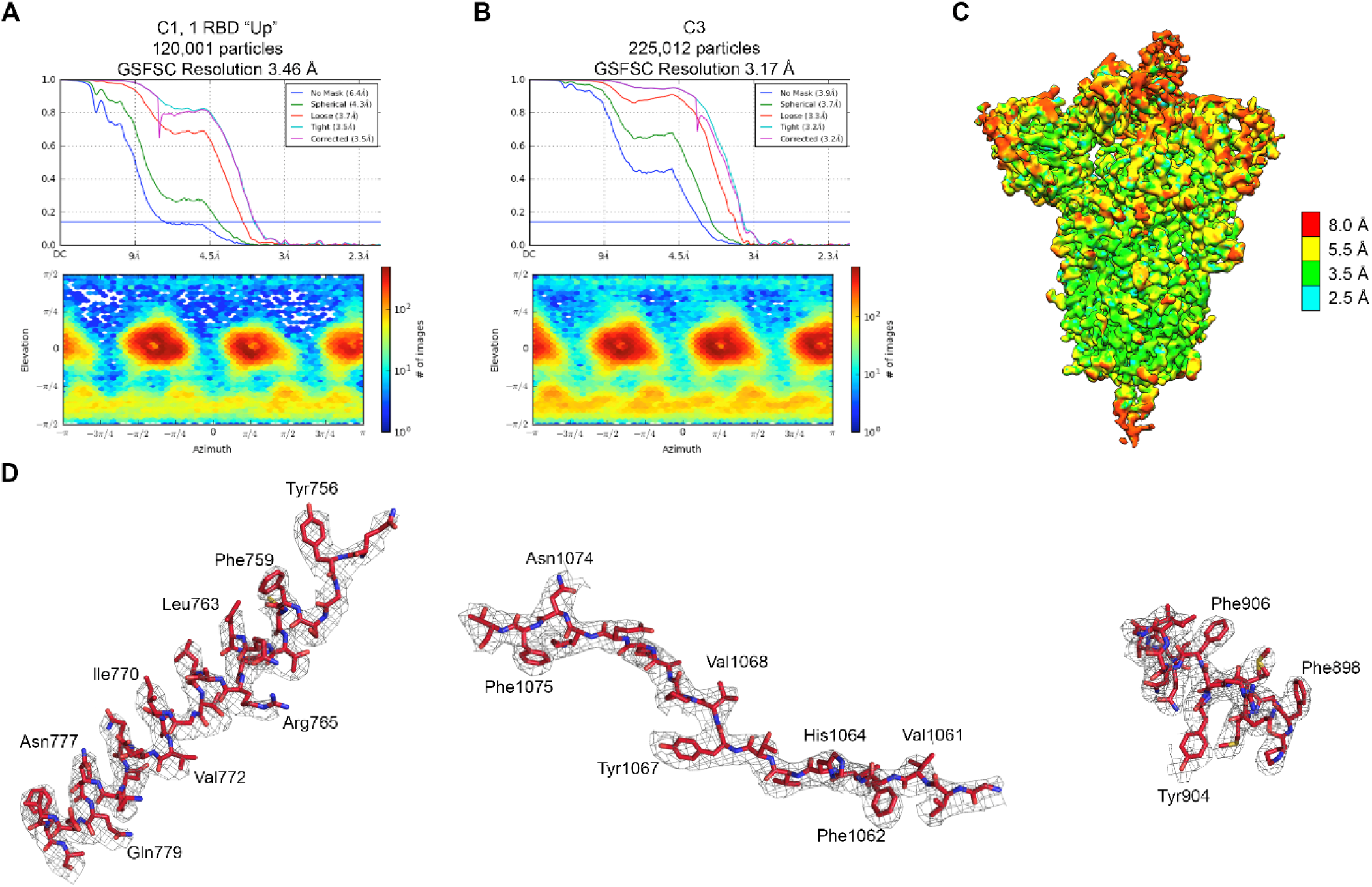
Cryo-EM structure validation. (**A**) FSC curves (*top*) and the viewing direction distribution plot (*bottom*) for 2019-nCoV S with a single RBD “up”. (**B**) FSC curves (*top*) and the viewing direction distribution plot (*bottom*) for the 2019-nCoV S processed with C3 symmetry. (**C**) The cryo-EM density of the 2019-nCoV S with a single RBD “up” is shown, colored according to local resolution. (**D**) Density from S2 of the C3-refined 2019-nCoV S structure. Residues are shown as sticks, colored according to **Figure 1A** with oxygen atoms colored red, nitrogens colored blue and sulfurs colored yellow. The cryo-EM density map is shown as a gray mesh.

**Supplementary Figure 3.**
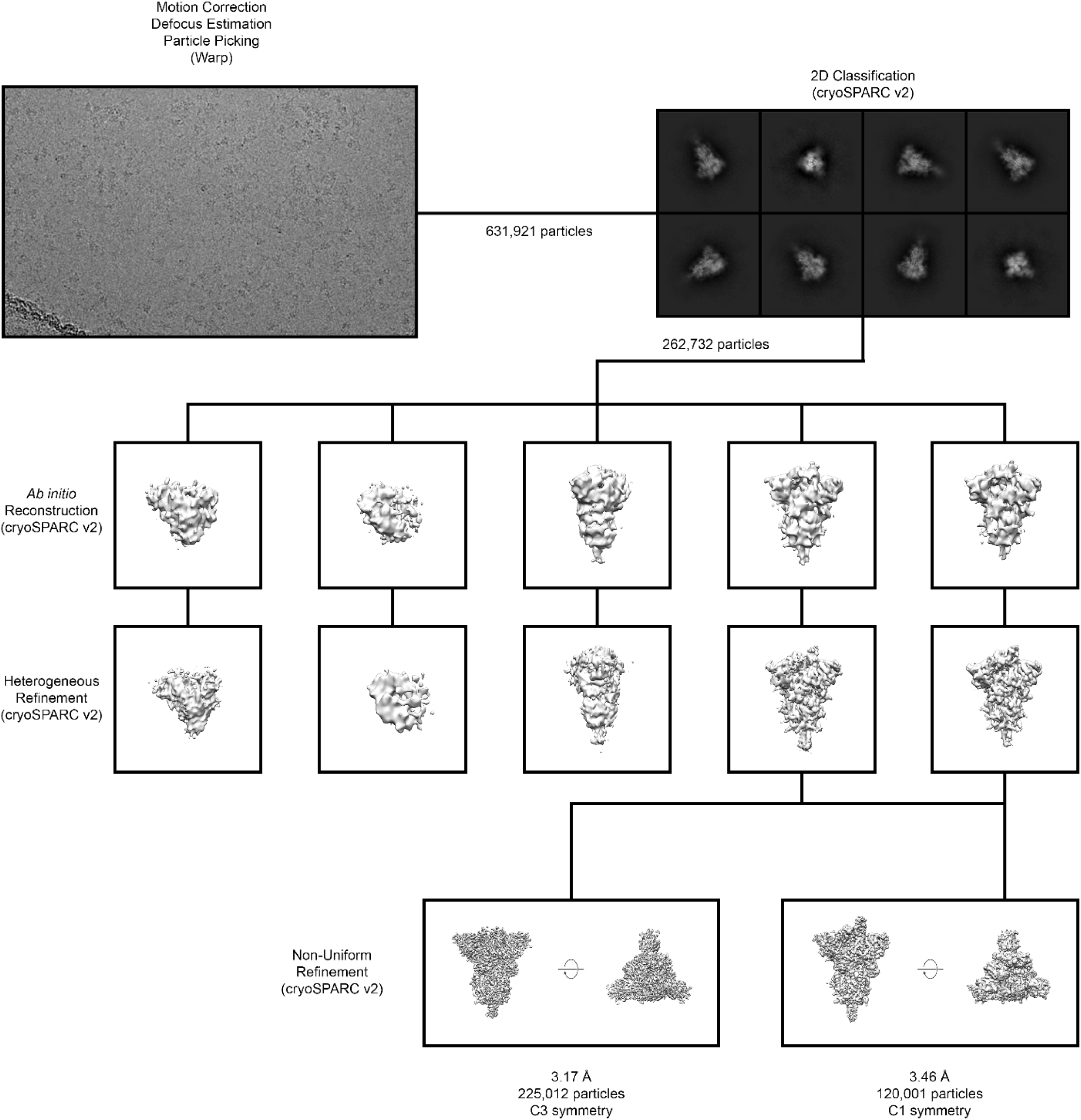
Cryo-EM data processing workflow.

**Supplementary Figure 4.**
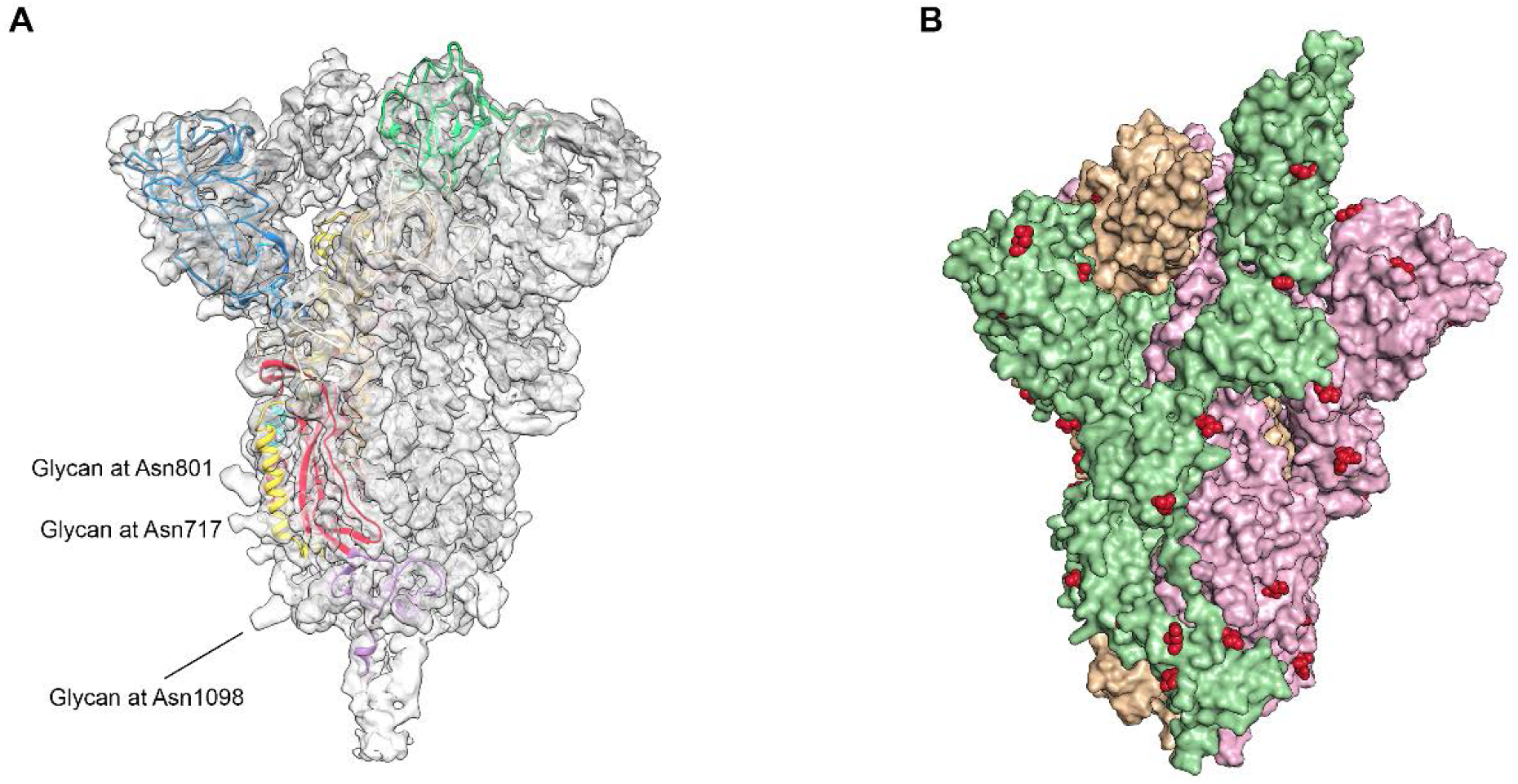
Cryo-EM map and *N*-linked glycosylation sites. (**A**) The unsharpened cryo-EM density map for the C3-processed 2019-nCoV S is shown as a transparent molecular surface, with a single protomer fit into the map shown in ribbons and colored according to **Figure 1A**. Some S2 density that corresponds to *N*-linked glycans is labeled. (**B**) The 2019-nCoV S trimer is shown as a molecular surface with each protomer colored green, pink or tan. Asparagine residues that correspond to *N*-linked glycosylation sites are shown as red spheres.

**Supplementary Figure 5.**
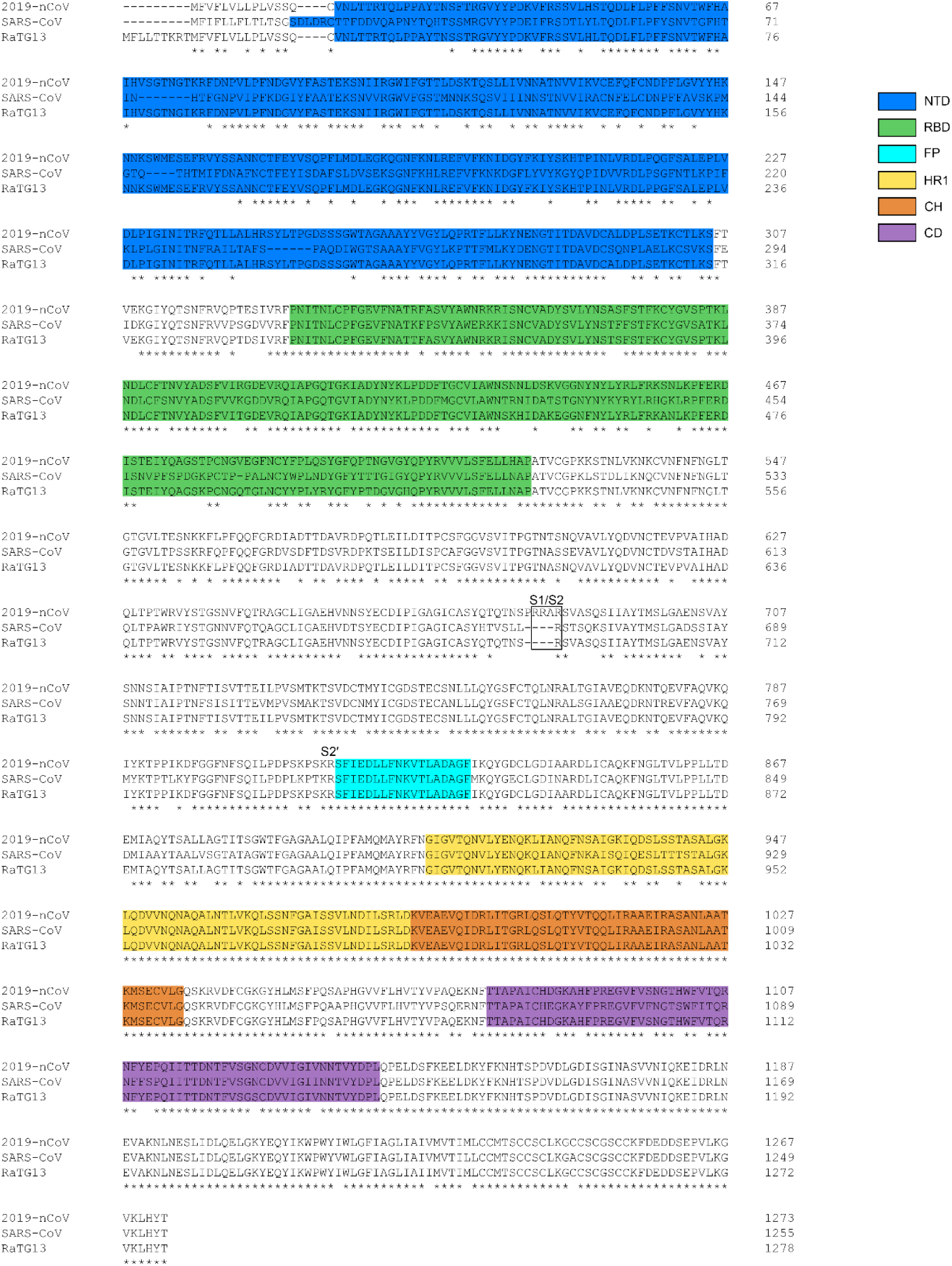
Sequence alignment of 2019-nCoV S, SARS-CoV S and RaTG13 S. Identical residues are denoted by an “*” beneath the consensus position. Structural domains are colored according to **Figure 1A**.

**Supplementary Figure 6.**
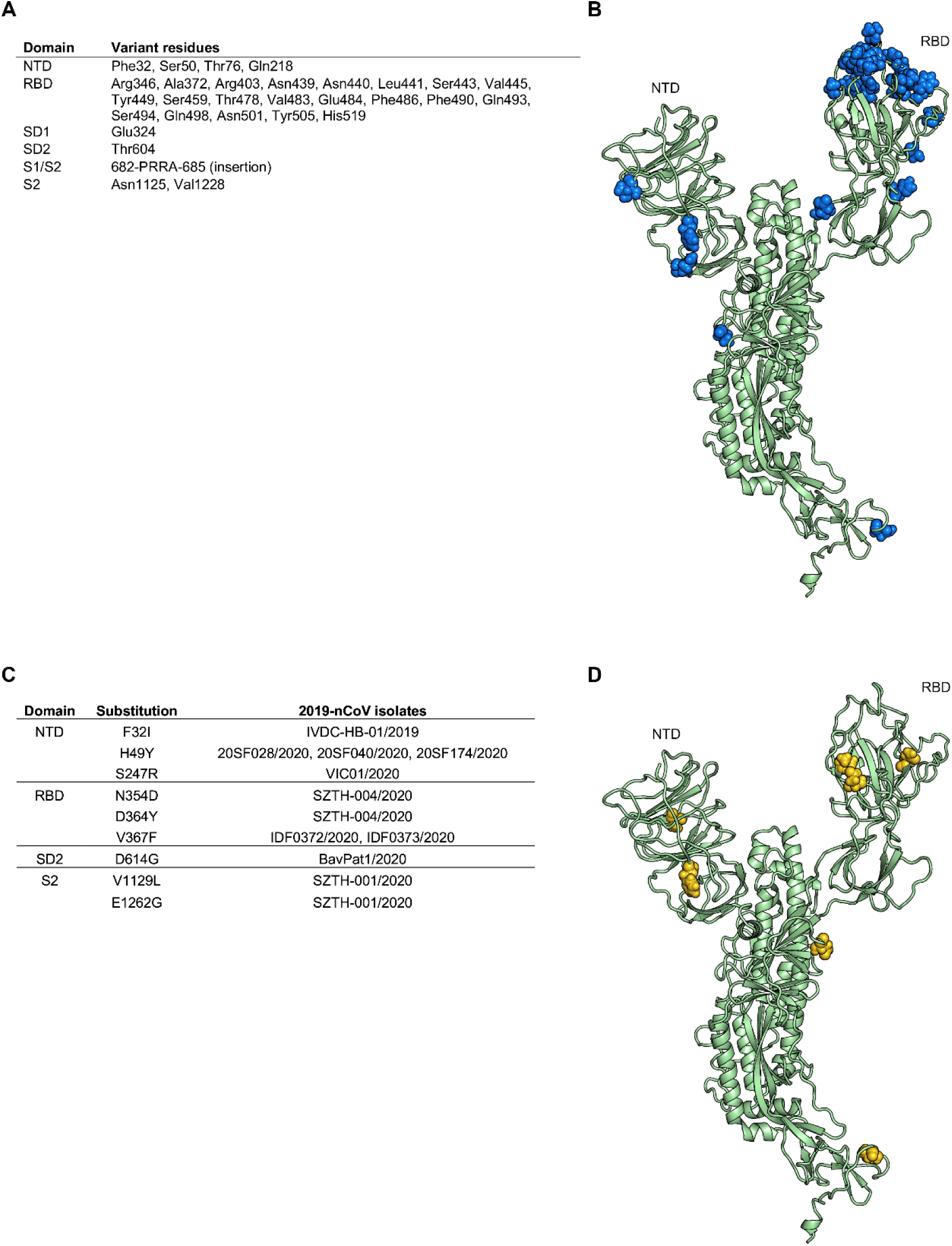
Sequence variability between RaTG13 S and 2019-nCoV S clinical isolates. (**A**) Table shows residues in the 2019-nCoV S protein that vary in RaTG13, grouped by structural domain. (**B**) A single monomer of the 2019-nCoV S protein is shown in ribbons, colored green. RaTG13 variant residues are shown as blue spheres. (**C**) Table shows variations in the 2019-nCoV S sequence based on 61 clinical isolates and the domains wherein these variations occur. (**D**) A single monomer of the 2019-nCoV S protein is shown in ribbons, colored green. Variant residues are shown as gold spheres.

**Supplementary Figure 7.**
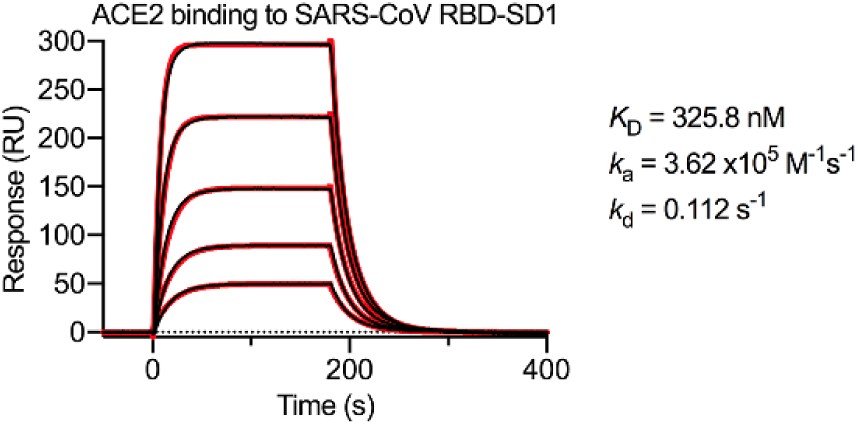
SARS-CoV RBD-SD1 binding to human ACE2. An SPR sensorgram is shown, displaying the binding between soluble human ACE2 and immobilized SARS-CoV RBD-SD1. The data are shown as black lines and the best fit of the data to a 1:1 binding model is shown in red.

**Supplementary Figure 8.**
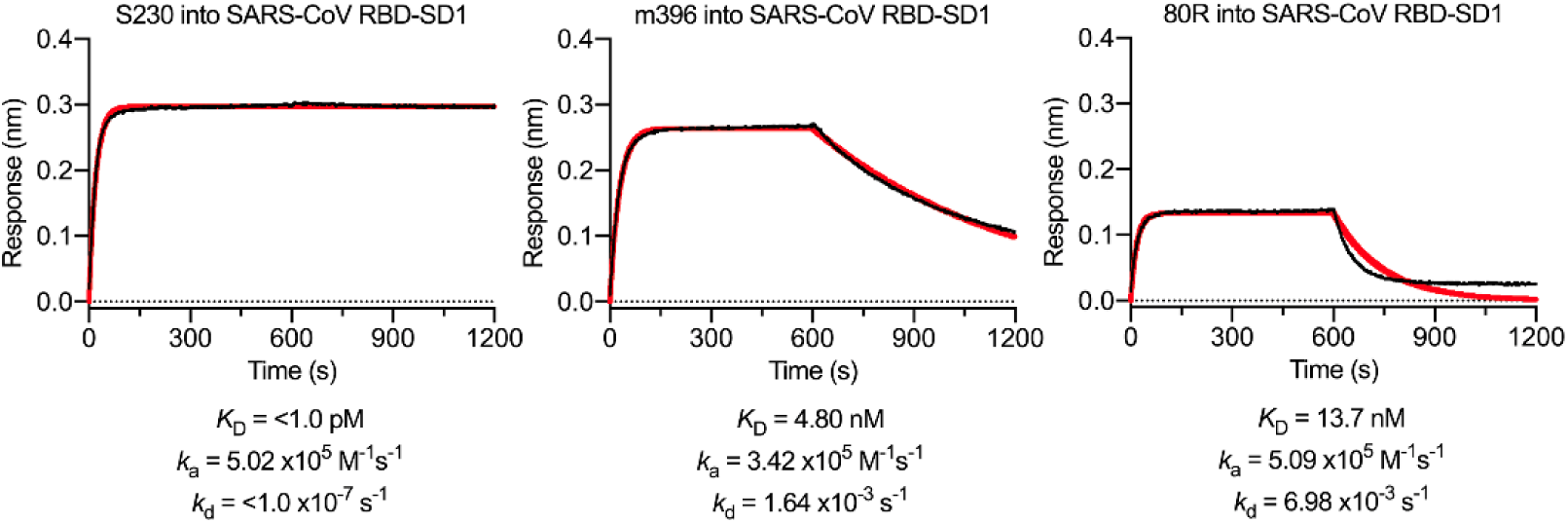
SARS-CoV RBD-directed antibody validation. The monoclonal antibodies that were tested for cross-reactivity to the 2019-nCoV RBD-SD1 were also tested for binding to the SARS-CoV S RBD-SD1 as a positive control. Binding data are shown as a black line and the best fit of the data to a 1:1 binding model is shown in red.

**Supplementary Table 1.**
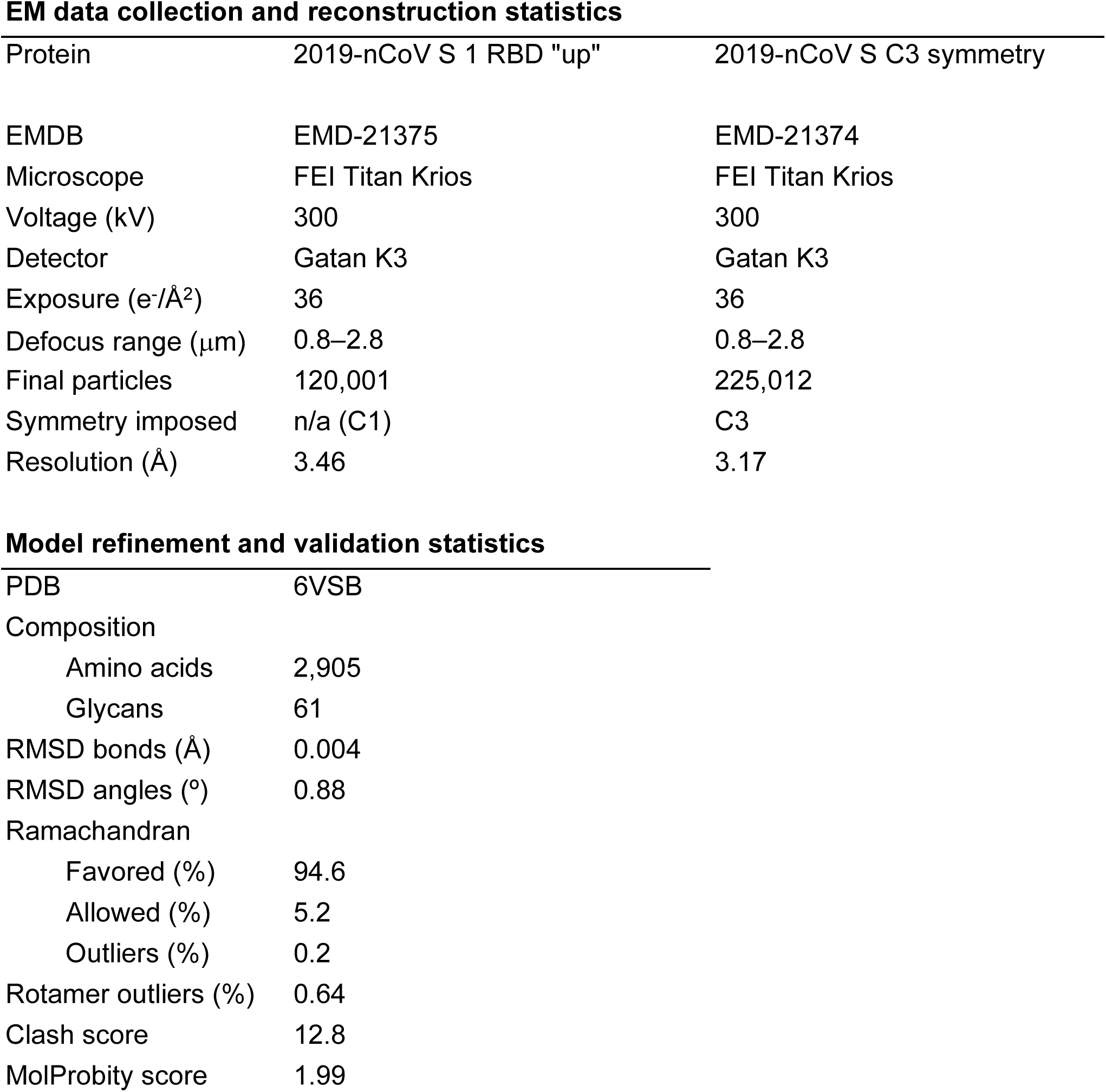
Cryo-EM data collection and refinement statistics.

**EM data collection and reconstruction statistics**
| Protein | 2019-nCoV S 1 RBD "up" | 2019-nCoV S C3 symmetry |
| --- | --- | --- |
| EMDB | EMD-21375 | EMD-21374 |
| Microscope | FEI Titan Krios | FEI Titan Krios |
| Voltage (kV) | 300 | 300 |
| Detector | Gatan K3 | Gatan K3 |
| Exposure (e<sup>-</sup>/Å<sup>2</sup>) | 36 | 36 |
| Defocus range (μm) | 0.8–2.8 | 0.8–2.8 |
| Final particles | 120,001 | 225,012 |
| Symmetry imposed | n/a (C1) | C3 |
| Resolution (Å) | 3.46 | 3.17 |

**Model refinement and validation statistics**
|  |  |
| --- | --- |
| PDB | 6VSB |
| Composition |  |
| Amino acids | 2,905 |
| Glycans | 61 |
| RMSD bonds (Å) | 0.004 |
| RMSD angles (°) | 0.88 |
| Ramachandran |  |
| Favored (%) | 94.6 |
| Allowed (%) | 5.2 |
| Outliers (%) | 0.2 |
| Rotamer outliers (%) | 0.64 |
| Clash score | 12.8 |
| MolProbity score | 1.99 |

**Supplementary Movie 1.** CryoSPARC 3D variability analysis. 2019-nCoV S trimer viewed from the side, along the viral membrane.

**Supplementary Movie 2.** CryoSPARC 3D variability analysis. 2019-nCoV S trimer viewed from the top, toward the viral membrane.

